# The Effect of Depriving the *Aedes aegypti* Mosquito of Natural Levels of Radiation

**DOI:** 10.64898/2026.06.29.735377

**Authors:** Liam Goodale, Cung Thawng, Immo Hansen, Geoffrey Smith

**Author notes:** Corresponding Author: (LG).

## Abstract

Organisms have spent their life histories exposed to background levels of natural ionizing radiation. To document the role that radiation plays, the deprivation of these natural levels has been studied by incubating organisms in the shielded space of underground laboratories. We report here on two studies (Study I and Study II) using *Aedes aegypti* for the first time as a model organism incubated 655 meters underground at the Waste Isolation Pilot Plant (WIPP) outside of Carlsbad, New Mexico, U.S.A. Male mosquitos were incubated at the surface exposed to natural background radiation, and were compared to two underground treatments in which incubators were supplemented with radiation sources used to mimic background and these groups were compared to the underground, radiation-deprived treatment. In Study I, the mosquitos incubated underground in the absence of natural radiation had higher levels of mortality compared to those incubated at the surface and PCA plots of the two transcriptomes were clearly differentiated. Study II was conducted the following year and the experiment was narrowed to include only the surface control and underground, radiation-deprived treatment which allowed for four biological replicates. Again, there was a higher level of mortality in the mosquitos grown underground compared mosquitos grown at the surface. Transcriptomes were not as clearly differentiated by PCA analysis and fecundity data were similar between the two groups. Functional analysis of transcriptomic DEGs from two independent studies suggested there are stress responses in radiation deprived mosquitoes. The absence of a secondary stressor in Study II is discussed as an explanation for the transcriptome differences in the two experiments.

## Introduction

There is now a significant body of literature on the biological effects of sub-background radiation that have been performed using several model organisms. In a landmark and impressively ambitious set of experiments, Planel et al. (1987) grew paramecia (*P. tetraurelia*) and cyanobacteria (*Synechococcus lividus*) under variable radiation dose rates, growing them underground in the Pyrenees (sub-background radiation), at the surface (background) and aboard the Soviet Salyut 6 spacecraft (above background radiation) [1]. In both model organisms, growth was inhibited in sub-background levels and, intriguingly, this inhibitory effect was abrogated when the cells were supplemented with radiation sources (^60^Co or ^232^Th) to mimic background levels of radiation. When the cells were exposed to levels of radiation about 20 times background during growth while in orbit aboard Salyut 6, both cell types proliferated at higher rates compared to background [1]. Planel’s pioneering work demonstrating the possibility of a fitness cost in depriving organisms of background radiation has certainly stimulated further research.

Satta et al. (1995) showed that *Saccharomyces cerevisiae* was more susceptible to DNA-damaging agents when grown in sub-background levels of radiation underground at the Italian Gran Sasso lab compared to surface-grown cells [2]. In more recent work at the Gran Sasso lab, Morciano et al (2018) showed evidence sub-background radiation reduced fertility in *Drosophila melanogaster* and, interestingly, viability was affected for several generations after being removed from the Gran Sasso underground [3]. In a follow-up experiment, Porrazzo et al. (2022) showed that incubation of *Drosophila* at Gran Sasso made flies more susceptible to DNA double-stranded breaks when exposed to ionizing radiation. Reminiscent of Planel’s work, when flies were grown underground in the presence of a supplemental gamma source, the radiation sensitivity was rescued. [4]

Mammalian tissue culture cells have an increased mutation rate [5] and be growth-inhibited after incubation in sub-background radiation [6]. In recent work at Canada’s SNOLAB, Lapointe et al. have elegantly simplified the biological model in their underground lab by incubating desiccated yeast cells long-term under sub-background conditions [7]. Rehydrating the yeast after 48 weeks, the group documented a significant inhibition in survival in cells incubated in sub-background conditions, and this reduced survival was accentuated in a DNA repair-deficient mutant [7]. Research on the biological effects of sub-background radiation is now on-going at 14 deep underground laboratories in 12 different countries [8].

In our work incubating organisms underground at the Waste Isolation Pilot Plant (WIPP) located near Carlsbad, NM, we have documented a stress response in bacteria [9,10,12], nematodes [13], and mammalian cells [11]. *Aedes aegypti* are well-studied insects and have been useful models in radiation biology [15]; we report in this paper the first use of the *Ae. aegypti* mosquito in sub-background radiation research. We have incubated male *Ae. aegypti* in the WIPP underground, and in one experiment we have separately “added back” two natural sources of radiation potassium chloride (KCl) and Pozzolana volcanic ash, to test for any rescue effects. Effects on mortality, reproductive success and gene expression (RNA Seq) are reported.

## Methods

### Experimental Overview

Two studies were performed at the WIPP facilities to investigate the effect on incubating *Ae. aegypti* (University of Georgia, UGAL, strain) in the presence of normal levels of background radiation compared to below-background radiation using methods we’ve developed with other biological models [10, 13]. In Study I, a single population of 200 male mosquitos were incubated in a control surface lab at WIPP and this population was compared to a population incubated underground in a pre-WWII steel vault to further decrease radiation exposure. In two other incubators underground, mosquitos were grown in the presence of two natural sources of radiation (KCl and Pozzolana volcanic ash), in an attempt to mimic natural radiation. In Study II, smaller populations of males (60) were incubated in four replicate smaller chambers for 15 days under only two conditions, at the surface and underground in the vault, and these were separately sampled for transcriptome analyses. In addition, after the 15-day incubation, mating assays were performed on the males, and egg counts and hatching success were quantified.

### Radiological Measurements

Table 1 shows the radiation dose rates obtained by repeated measurements of at least 15 hours using a gamma-specific ion chamber (Fluka model 451P, calibrated yearly) from the surface and underground at WIPP. The levels of radiation in the underground vault are below instrumental detection limits [10], and so we have performed a Monte Carlo Neutral Particle (MCNP) estimation (ca. 0.004 nGy/hr) and added that to the maximum radon concentration measured in the underground at WIPP (0.004 nGy/hr) [11] to give us the estimated dose rate of 0.01 nGy/hr shown in Table 1. In Study I, following the lead of Planel et al. (1987) [1], we have compared our radiation-deprived treatment to underground controls where we have supplemented the radiation to mimic surface levels in which we have grown organisms underground in incubators where we have “added back” kg quantities of KCl as a natural source of radiation [10,11,12].

**Table 1:**
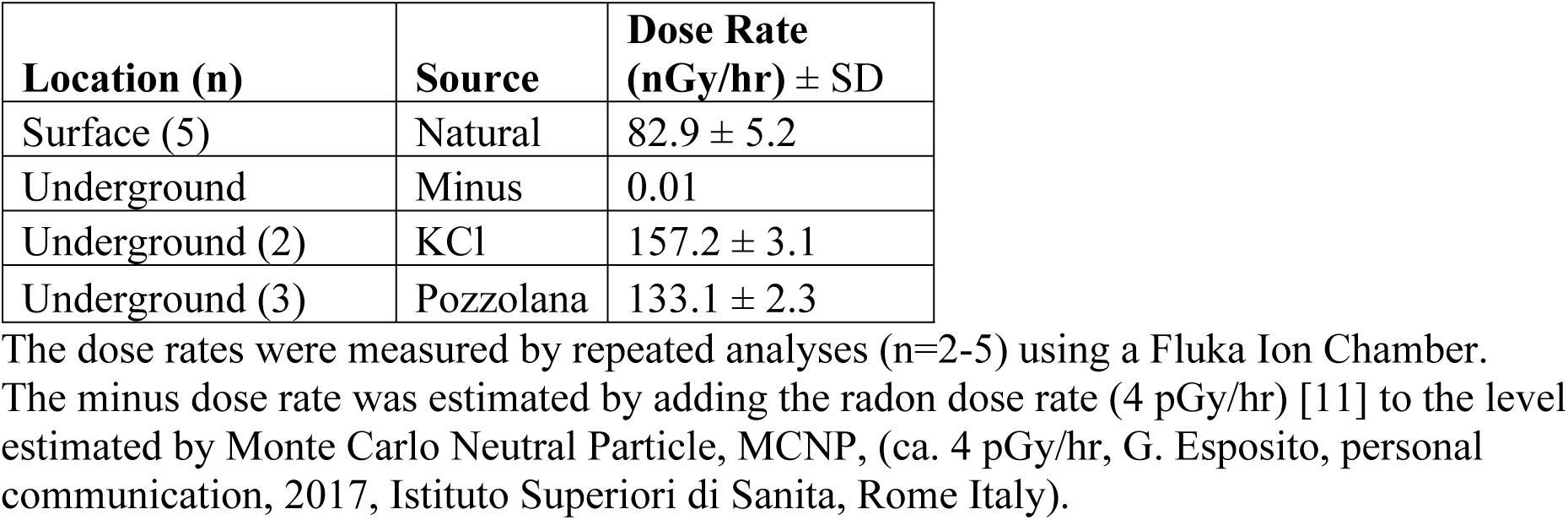
Gamma Dose Rates Used in this Study.

| **Location (n)** | **Source** | **Dose Rate (nGy/hr) ± SD** |
| --- | --- | --- |
| Surface (5) | Natural | $82.9 \pm 5.2$ |
| Underground | Minus | 0.01 |
| Underground (2) | KCl | $157.2 \pm 3.1$ |
| Underground (3) | Pozzolana | $133.1 \pm 2.3$ |

The dose rates were measured by repeated analyses (n=2-5) using a Fluka Ion Chamber. The minus dose rate was estimated by adding the radon dose rate (4 pGy/hr) [11] to the level estimated by Monte Carlo Neutral Particle, MCNP, (ca. 4 pGy/hr, G. Esposito, personal communication, 2017, Istituto Superiori di Sanita, Rome Italy).

One goal of the current research was to test another, potentially more natural radiation source for our add-back controls, and that is a volcanic tuff-based material called Pozzolana. In terms of quantities of radiation added back, Pozzolana (133 nGy/hr) provided a similar dose rate as KCl (157 nGy/hr), and both are representative of typical surface level radiation (114-570 nGy/hr) [16]. See Table 1 for all pertinent dose rates.

### Mosquito Incubation Conditions and Post-incubation Mating Assay

Mosquito hatching and rearing was conducted at New Mexico State University (NMSU) using methods previously published in the Hansen lab [17]. Males and females were segregated within 24 hours of emerging and were assumed to be virginal [18, 19]. At NMSU as part of Study I, a population of 210 male mosquitoes were placed in a large (31 cm x 31 cm) Bug Dorm (Megaview Science, Taiwan), and four dorms were used in the four experimental groups (Surface, Minus, KCl, and Pozzolana, fed 20% sucrose), and these were transported to WIPP, four hours away. In Study II, eight small Bug Dorms (11 cm x 11 cm) were prepared with 60 males each. These would become the Minus and Surface treatment groups with four replicates each (M1, M2, M3, M4, S1, S2, S3, and S4, fed 10% sucrose). The dorms were then inserted in the Surface and Vault incubators, but for the radiation-amended treatments (KCl or Pozzolana), the dorms were inserted in irradiators made with 6 panels that surrounded the bug dorms. The panels were either filled with quantities of KCl (2.3 kg/panel) or with Pozzolana (2.1 kg/panel) estimated to give the same dose rates. **Figure 1** illustrates the study site location, the World War II steel vault in the underground, as well as a large mosquito habitat inside an irradiator. All incubators at WIPP were maintained between 22.7-24.3 °C, 75-95% RH and were under a 14-hr light/10-hr dark cycle using LED strips as light sources (DZF Tech, China). It should be noted that in Study I, 40 mosquitos escaped in the underground during the first 5 days of incubation, so the surface population was culled to match the underground population. As a result, both populations were 162 +/- 5 on day 5.

**Figure 1:**
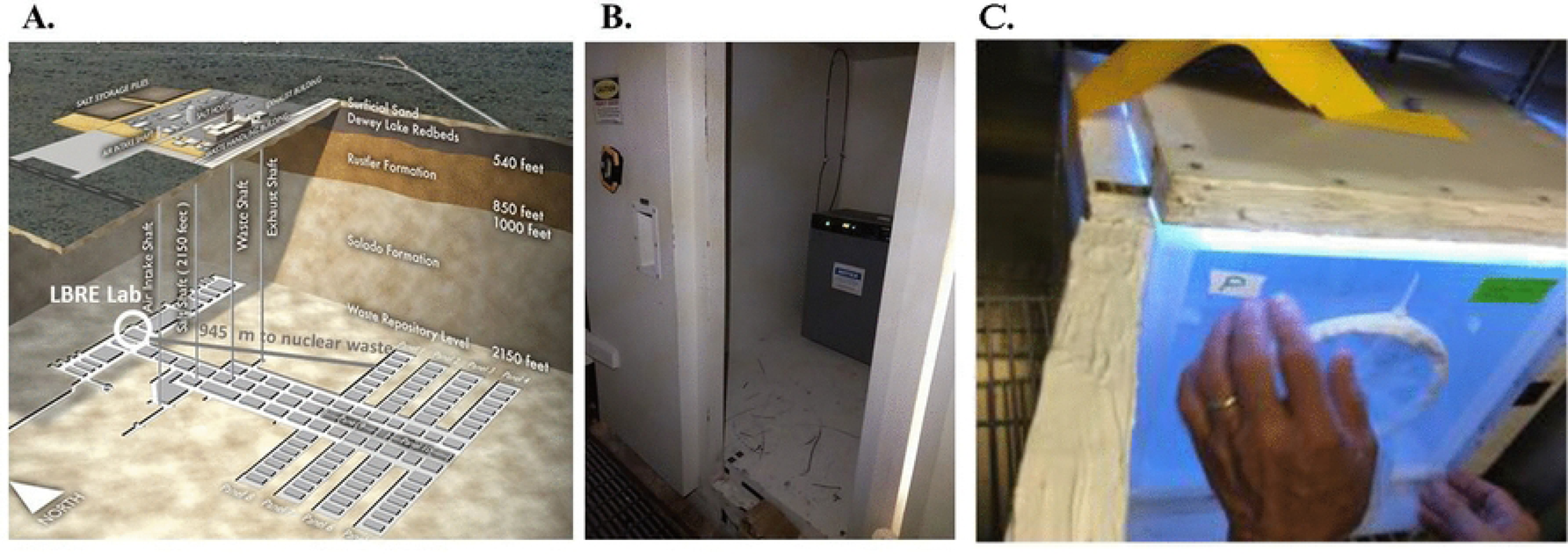
Location of Study. A. Location of the Low Background Radiation Experiment (LBRE) in the WIPP mine. B. An incubator in the Pre-World War II vault underground at WIPP. C. Pozzolana control group, with irradiator opened for observation, underground at WIPP.

In Study II, on day 15 of the incubations, a mating efficacy test was performed on 40 males from each of the four replicates of the Minus and Surface treatments, with the males being paired with females that had emerged from the pupal stage 2 - 4 days previously. In preparatory steps at NMSU, females were added to a perforated 50 mL plastic conical tube having a moistened paper disk, and for each treatment, 20 tubes having one female and 20 tubes having two females were prepared and transported to WIPP. Males were aspirated from their bug dorms and were anesthetized on ice and added to the mating chambers with females that had been gassed with CO_2_ for about three seconds. The mating tests for each incubation treatment replicate consisted of two ratios of males to females, 1:1 and 1:2. Mating was carried out overnight (20 +/- 2 hrs) in incubators set at 24°C and 90% relative humidity.

After the overnight mating assay, females were exposed to CO_2_ for three seconds in these tubes and were transferred to 16 separate small bug dorms to keep the ratios and treatment replicates separate and they were transported to NMSU with access to 10% sucrose water. After two days, blood feeding was initiated under observation for approximately 7 hours. Afterwards, the females were returned to their egg laying chambers and the tubes were partially submerged in water to promote ovipositing. Based on the method described by Kandel et al. [18], egg laying was allowed for 6 days and then the papers were collected and the egg counts tabulated.

### RNA Extractions and Transcriptome Software Comparisons

In both Study I and Study II, RNA extractions were carried out using the following procedure: mosquitoes were aspirated and anaesthetized on ice and triplicate samples of 20 mosquitoes were placed in a 1.8 mL microfuge tube having 0.5 mL Qiagen RNeasy lysis buffer and were homogenized 30 sec. with a battery-operated, vibrating pestle. Samples were immediately frozen, transported to NMSU and the RNA extractions were completed the following day using an RNA isolation kit (RNAeasy Mini Kit, QIAGEN) according to manufacturer’s instruction. Approximately 2 micrograms of frozen RNA was shipped overnight to Azenta for sequencing and bioinformatic analysis.

RNA-Seq raw reads into ArrayStar™ along with the *Aedes aegypti* Liverpool strain AGWG (AaegL5.0) genome FASTA (.fna) and annotation (.gbff) files as the reference genome [20]. Two software packages (CLC and Azenta) were used to analyze the transcriptome data. The CLC Genomic Workbench 22.2 software package uses a Local alignment to map sequences to a reference genome, the trimmed mean of M-values (TMM) to normalize and its own CLC DEG statistical package [21]. The Azenta software uses the STAR aligner to map, the median-f-ratios to normalize and the DESeq2 for statistical analyses [22].

### Gene Ontology Term Analysis

Gene ontology (GO) terms are used in many cases to summarize complex genetic profiles, and can be predictive of collective function [23, 24]. GO terms used to describe DEGs herein were provided by the DAVID bioinformatics tool [25] as well as VectorBase [26].

## Results and Discussion

### Radiological Conditions

The U.S. Environmental Protection Agency estimates the U.S. natural background radiation to be 410 nGy/hr, but it varies by location with the highest average dose in South Dakota at 1100 nGy/hr and the lowest dose in Florida at 150 nGy/hr [27]. Gamma radiation is commonly measured in sub-background radiation research [5,11] due to its accuracy of measurement and gamma being an important component of natural background, representing approximately 25% of natural sources [28]. In this study, the basic comparison is between natural background radiation at the WIPP surface (82.9 nGy/hr gamma) and the abnormally low radiation (ca. 0.01 nGy/hr) that penetrates the 655 meters of rock and salt overburden and the 15.2 cm-thick pre-WWII steel vault located in our experimental area underground (**Table 1**). In Study I, underground incubators were supplemented with a new source of natural radiation (Pozzolana, a volcanic fly ash) and this was compared to the KCl-radiation amendment that we’ve used previously as an underground “radiation add-back” control [11,12].

An advantage of using mosquitos as a biological model in sub-background radiation experiments is that the nutritive requirements for adult males are much simpler than other organisms used to date in sub-background radiation work, that is, they only require sucrose [18]. Lampe et al. [29] reported that in below background radiation studies, the most important contributor to test organisms’ internal radiation in shielded underground laboratories is the ^40^K in the feedstock.

Lampe et al. also state that the contribution from “^14^C is negligible,” which is consistent with its occurrence in nature: there are only two ^14^C atoms in 10^12^ non-radioactive carbon atoms, in contrast to ^40^K which occurs once in 10^4^ potassium atoms [30]. Additionally, compared to our previous work when using bacteria [10, 12, 14], nematodes [13] and mammalian tissue culture cells [11], we have removed all other external sources of radiation from our underground vault. And so, the results reported here from the below background radiation treatment are significantly lower (approximately 100 times) than we’ve previously reported.

### Male Mortality and Mating Success

Pilot studies showed that males of the *Ae. aegypti* Liverpool strain grown at NMSU experienced more than 50% mortality within two weeks, whereas the UGAL strain typically exhibited 20% mortality, and so the UGAL strain was chosen for this study [31]. In Study I, 200 one to three-day-old males were distributed in three large BugDorms, one was kept at NMSU, and two others were transported (ca. four hours at 22-25 °C) and grown at the WIPP surface or in the vault underground. Over the first 12 days, the UGAL strain exhibited a mortality rate of 5-10%, but increased to 15-20% in four days (**Figure 2A**). It is important to note that the death phase also occurred in mosquito populations that were kept at the surface 230 miles away at NMSU; equally important is the observation that both controls declined 15% but the underground treatment mosquitos declined 20% over the last four days of incubation. However, since there was only a single population of 200 at each location, the trend of increased mortality in the WIPP underground was not statistically significant.

**Figure 2:**
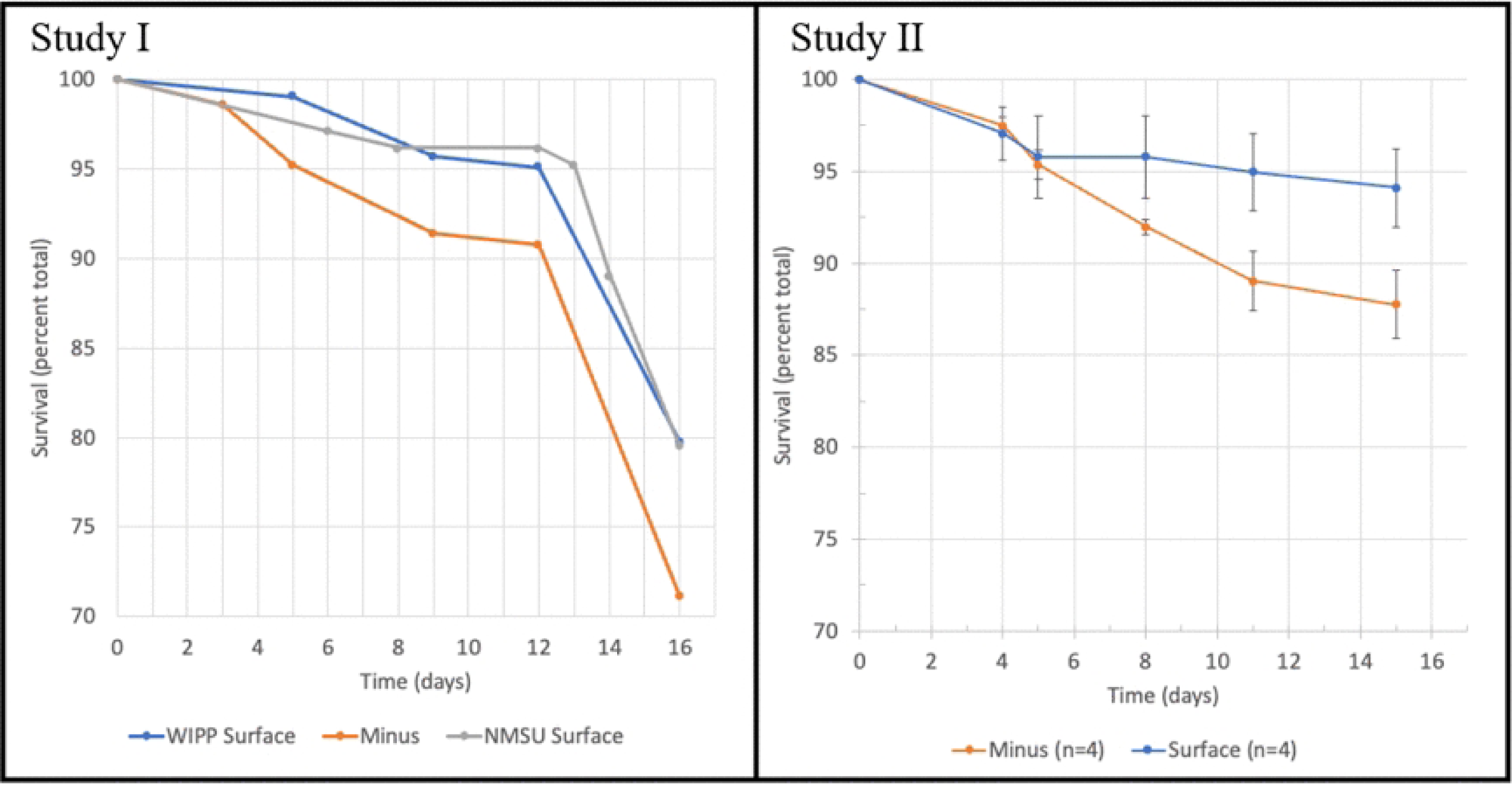
Mortality plots from Study I (A) and Study II (B) studies. Study I had a single BugDorm at the Surface and Underground (n ∼200 mosquitos/dorm) and Study II had four replicate BugDorms at each location (n ∼ 60/dorm). Kaplan-Meier analysis of the data shown in 2B showed the sub-background radiation to have a higher mortality than surface levels (p=0.002).

A simplified version of this first study was repeated the next year (except with 10% sucrose instead of 20%) using four replicate, smaller BugDorms with 60 mosquitos per dorm. Similar to the first study, the mortality rates were stable for the first 12 days, but Study II had to be terminated at day 15 due to a problem at WIPP. There was a higher mortality rate in the underground-grown mosquitos, with respective 6% and 12% mortality rates for the WIPP surface and underground mosquitos at day 15 (**Figure 2B**). Using the Kaplan-Meier estimator it was found that the mortality rates between mosquitos grown underground were significantly higher than when grown at the surface (p=0.002). Interestingly, in the first 12-13 days, the kinetics of the mortality rates of Study I and Study II were almost exactly alike, with the surface control dying off at half the rate of the radiation-deprived underground treatment. It is unfortunate that we had to terminate Study II early before the higher mortality phase, that was documented in the first study, had ensued.

In Study II, after the two-week incubations at WIPP, approximately 20 replicate males were mated with 3-day-old females in female:male ratios of 1:1 or 2:1. The intention in using a male paired with two females was to challenge the males and perhaps accentuate any radiation treatment differences. Though there was a trend for less mating success and a lower number of eggs laid from the 2:1 ratio, the differences were non-significant (**Table 2**). There was no statistical difference between the two radiation groups of males in terms of mating success and resultant number of eggs produced. The percentage of eggs that hatched was also not statistically different between the two treatments (data not shown).

**Table 2.**
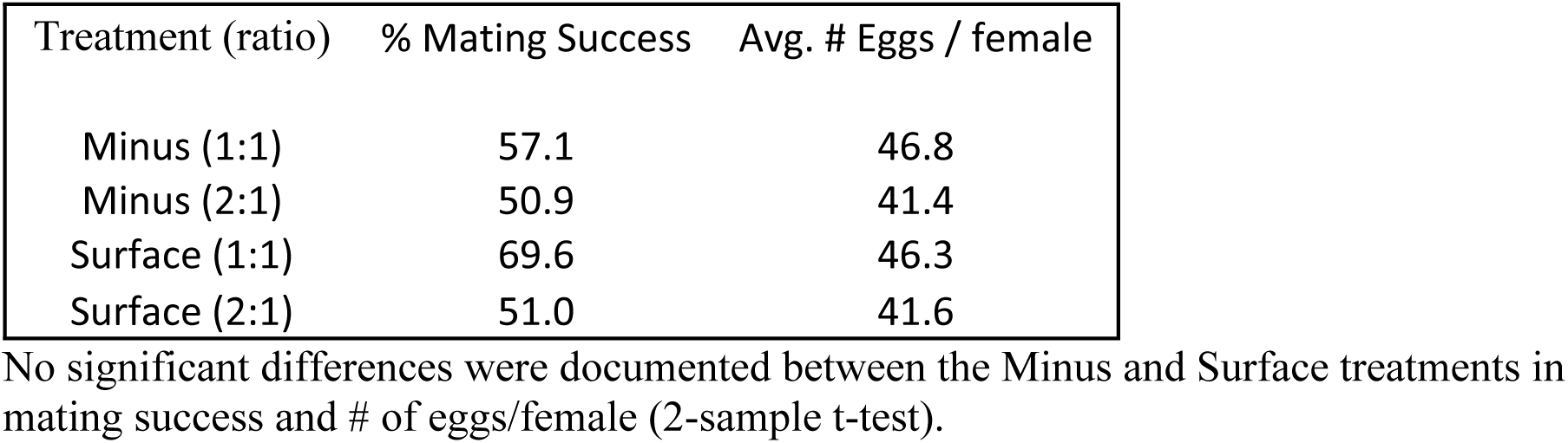
Mating success and egg number. In the Study II, after incubations at WIPP, males were mated with females and the percent of females which laid eggs (% mating success) and the average number of eggs laid is shown.

### Transcriptional Analyses

In Study I, 20 mosquitos of one population of mosquitos from each location was subsampled three times to provide three replicate transcriptomes, and in Study II, 20 mosquitos from four separate populations were sampled from the two locations. We performed principal component analyses (PCA) using CLC (**Figure 3 A**) and Azenta (**Figure 3 B**) software. Consistent with previous results in which we’ve documented statistical gene expression differences between organisms grown in radiation sufficient and deficient conditions [10,11,12,13], the PCA analyses of the Study I transcriptome showed a clear separation between the expression patterns in mosquitos grown in control levels of radiation at the surface compared to those grown underground at WIPP. In Study II, the PCA patterns were similar but overlapped to some extent between the two conditions (**Figure 3**).

**Figure 3.**
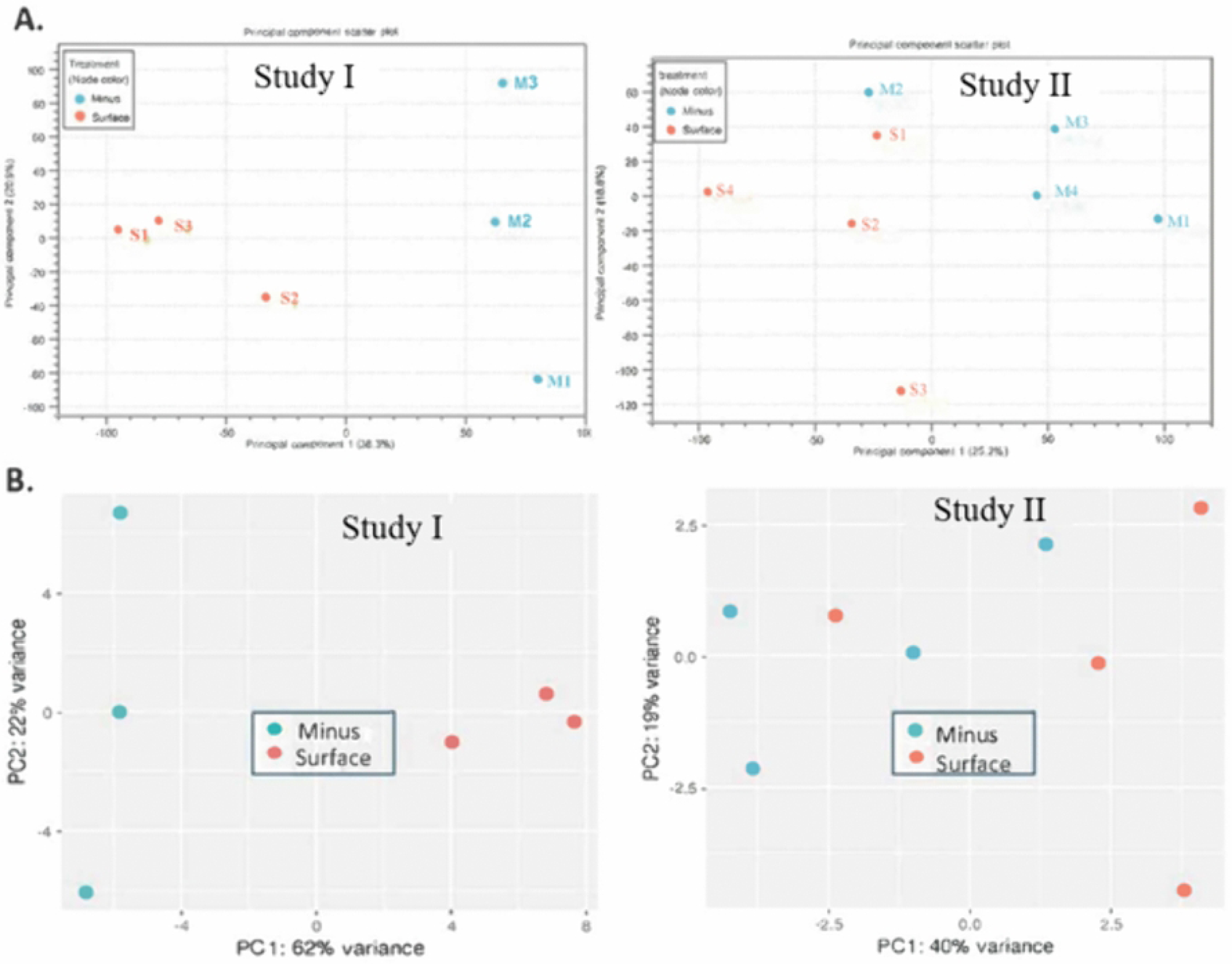
Principal component analyses by CLC (A) and Azenta (B) of the transcriptome data from Study I and II.

When differentially expressed genes (DEGs) were compared between both studies using Azenta software, the results were more contrasting, with 221 DEGs documented in Study I, but only seven identified in Study II (**Supplemental Table 1**). A parallel analysis using a different software package we’ve shown to be more conservative than Azenta (DNAStar’s DeSeq-2 pipeline) [22] resulted in similar ratios of DEGs (data not shown).

As we’ve discussed, in Study I, one large population was used in each treatment and in Study II, four independent populations were used. The Study II data would be expected to be more statistically valid, but the completely different sets of DEGs between the two years would indicate that the mosquitos were experiencing different conditions.

We propose that the explanation that largely accounts for these contrasting transcriptomes is based on the contrast in mortality rates in the two years, particularly in the last phase of the incubation. In Study I, the mosquitos had started to enter the stress phase of their life cycle, whereas in Study II, the mosquitos at the time of RNA harvest were still healthy, as is shown in the contrasting mortality curves in **Figure 2**. This growth phase-dependent response to below-background radiation is similar to what we’ve documented before in the bacterium *Shewanella oneidensis* in which, under the stress of reaching stationary phase, *S. oneidensis* exhibited a much stronger transcriptional response to radiation deprivation than when growing rapidly in early log phase [11]. Similarly, in a study by Muturi et al. [32], when larvae of *Ae. aegypti* were stressed by elevated temperatures or starvation, adults were more susceptible to a second stress of pathogen infection. The exposure of one stress that accentuates a secondary stress has been called stress sensitization [33] and may well explain the accentuated stress response in Study I. The GO terms falling under Molecular Function groups of DEGs identified in the two studies are shown in **Table 3**. As discussed above, there were only seven DEGs identified from Study II. Of the 33 upregulated genes from Study I, only four were assigned GO Molecular Function terms and these were associated with protein folding-chaperones. The main groups identified to be downregulated in Study I were overlapping groups of oxidoreductase and transition metal binding genes (totaling 59) and 25 genes identified as protease genes. We have similarly shown in bacteria oxidoreductase genes [11] and peptidases [12] to be among the more responsive groups of genes to the deprivation of natural levels of radiation.

**Table 3.**
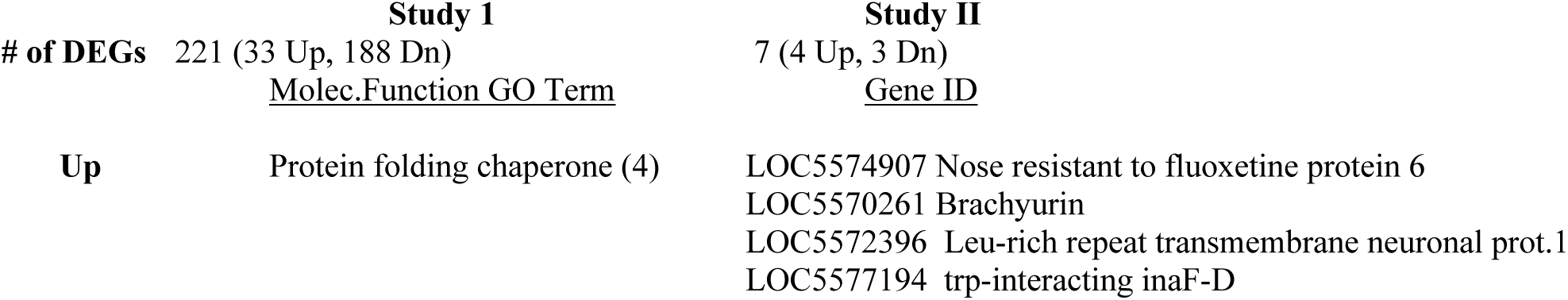
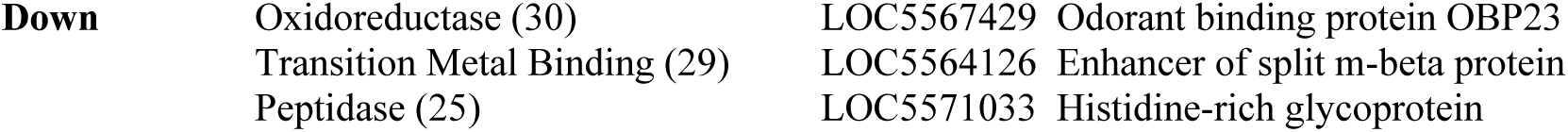
The number of differentially expressed genes (DEGs) are shown from the surface control and underground treatment represented by Molecular Function terms (Study I) and gene ID (Study II). Only genes that met the criteria of ≥ 2-fold change, FDR ≤ 0.05 and ≥ 30 reads are shown. Of the 33 upregulated genes from Study I, only 4 were assigned Molecular Function GO terms.

There were no conserved DEGs between both Study I and Study II meeting the criteria used herein, however, when lowering the threshold to 1.5x fold change for Study II, there are some conserved and functionally related DEGs. However, there were functionally similar or related DEGs that appeared in each experiment (**Table 4**). For example, multiple cytochrome P450s were upregulated in the Minus group each year (11 in Study I, and 2 in Study I). In addition, zinc finger proteins, myosin 3 proteins, and odorant binding proteins were also consistently downregulated.

**Table 4.** Related Downregulated DEG Comparison Between Experiments.

| **Study II (1.5-2-fold change)** |  | **Study I (2-fold change)** |  |
| --- | --- | --- | --- |
| **Gene ID** | **Name** | **Gene ID** | **Name** |
| **LOC5567429 (2x)** | General odorant-binding protein 68 | LOC5564586 | General odorant binding protein 99b |
| **LOC5569477 (1.5x)** | Zinc finger protein 160 | LOC5571252 | Zinc Finger Protein 664 |
| **LOC5571037 (1.5x)** | Chymotrypsin-1 | LOC5571039 | Chymotrypsin-1 |
| **LOC5571543 (1.5x)** | Cytochrome P450 6d3 | LOC5564852 | ninaC |
| **LOC5572380 (1.96x)** | Cytochrome P450 307a1 | 11 Cytochrome P450s | Refer to Table 5 |
| **LOC5572723 (1.5x)** | Myosin-IIIb |  |  |

The cytochrome proteins (CYPs) in each family are understood to share common function in *Aedes aegypti*. For example, the CYP9, CYP6, and CYP28 subfamilies of the CYP3 clan are associated with insecticide resistance, with subtle differences between them. Of the seven regulated genes of the CYP3 clan listed in **Table 5**, four are in the CYP9 subfamily, 2 in the CYP6 subfamily, and 1 is in the CYP28 subfamily. The CYP4 clan is associated with endogenous metabolism, whereas the CYP3 clan is associated with exogenous metabolism (especially xenobiotic toxins, such as insecticides). Three of the remaining 11 DEGs are in the CYP4 clade and last belongs to the Mitochondrial clade [34]. There is evidence that CYP3-clan (especially CYP6 and CYP9) belong to a stress-responsive detoxification system [35]. Studies have found that they are highly expressed when mosquitoes are exposed to natural (e.g., viruses) and artificial (e.g., insecticides) stressors [36, 37, 38].

**Table 5.** Study I CYP450 Downregulated DEGs.

| **Gene ID** | **Name** | **Clade** |
| --- | --- | --- |
| **LOC5578181** | Cytochrome P450 4C1 | 4 |
| **LOC5572942** | Cytochrome P450 4D2 | 4 |
| **LOC5571543** | Cytochrome P450 6D3 | 3 |
| **LOC5564751** | Cytochrome P450 9E2/9J28 | 3 |
| **LOC110677901** | Cytochrome P450 24A shade | Mitochondrial |
| **LOC5568653** | Probable Cytochrome P450 28A5 | 3 |
| **LOC5579144** | Probable Cytochrome P450 4AC1 | 4 |
| **LOC5565578** | Probable Cytochrome P450 6A14 | 3 |
| **LOC5564748** | Probable Cytochrome P450 9F2 | 3 |
| **LOC5564749** | Probable Cytochrome P450 9F2 | 3 |
| **LOC5571141** | Probable Cytochrome P450 9F2 | 3 |

There was no conservation of statistically significant upregulated DEGs between Study I and Study II that met our criteria (≥30 reads, ≥2-fold change in expression). However, there were functionally similar or related DEGs that appeared in each experiment (**Table 6**). For example, protein lethal (2) essential for life was consistently upregulated, and so too were multiple proteases. In Study I, there were also multiple heat shock proteins. Together, these results show evidence that there was a significant stress response in the radiation-deprived groups of mosquitos.

**Table 6.**
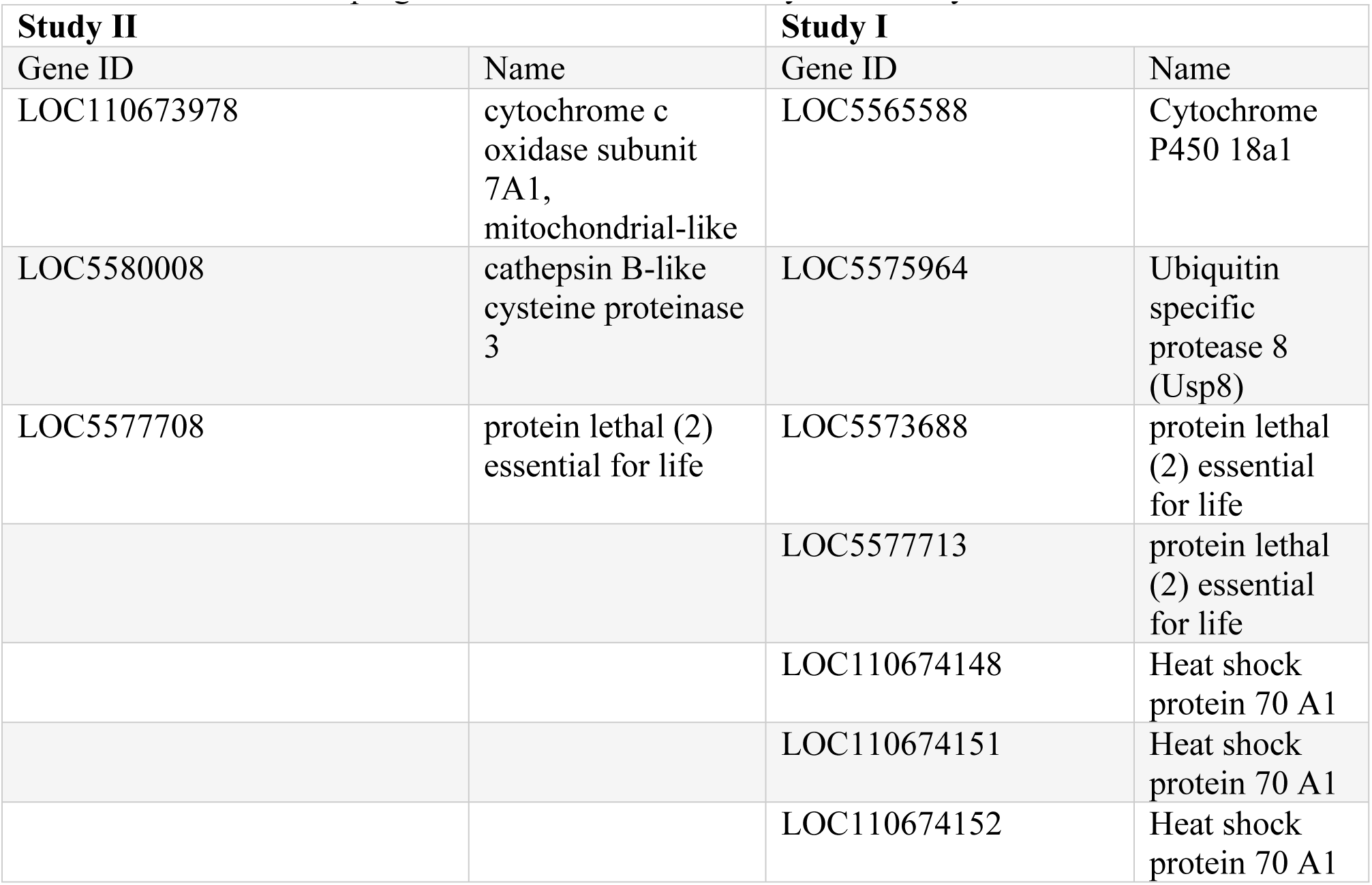
Some Related Upregulated DEGs Between Study I and Study II.

| **Study II** |  | **Study I** |  |
| --- | --- | --- | --- |
| Gene ID | Name | Gene ID | Name |
| LOC110673978 | cytochrome c oxidase subunit 7A1, mitochondrial-like | LOC5565588 | Cytochrome P450 18a1 |
| LOC5580008 | cathepsin B-like cysteine proteinase 3 | LOC5575964 | Ubiquitin specific protease 8 (Usp8) |
| LOC5577708 | protein lethal (2) essential for life | LOC5573688 | protein lethal (2) essential for life |
|  |  | LOC5577713 | protein lethal (2) essential for life |
|  |  | LOC110674148 | Heat shock protein 70 A1 |
|  |  | LOC110674151 | Heat shock protein 70 A1 |
|  |  | LOC110674152 | Heat shock protein 70 A1 |

### Supplemental Radiation Sources to Mimic Background

In some of our studies on the effects of natural background radiation we have used potassium as natural source of radiation since it is the most important terrestrial source of radiation [30] and, due its Compton scattering, it represents a broad spectrum of ionizing energies [10]. In collaboration with G. Esposito at Rome’s Istituto Superiori di Sanita [39], we implemented a volcanic source of radiation, a type of fly ash called Pozzolana. In addition to having K-40, pozzolana also has uranium and thorium, two other important sources of terrestrial radiation [40]. At Day 16 at the end of the incubation, 28.8% of the 200 mosquitos had died in the underground treatment, 20.2% had died at the Surface and 21.7% and 13.7% had died underground with KCl and Pozzolana supplements, respectively (**Figure 4**). Though not statistically different, the mosquitos underground supplemented with two sources of natural radiation (KCl or Pozzolana) appeared to be rescued back to surface control levels compared to the unsupplemented, sub-background treatment.

**Figure 4.**
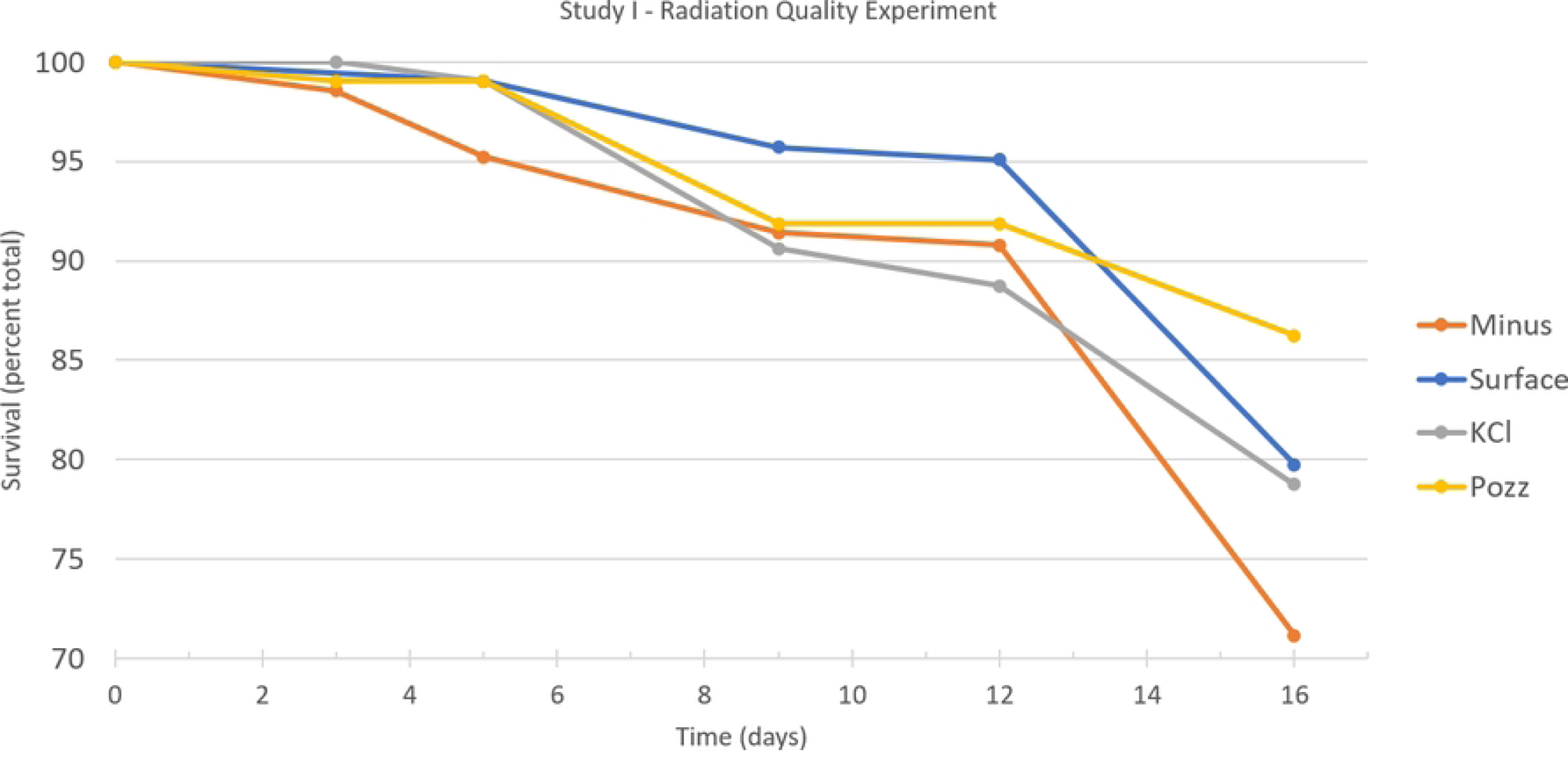
Mosquito mortality as affected by radiation source. Four sets of 200 mosquitos were grown at the Surface or the WIPP underground: “Minus” mosquitos were grown in the vault and two underground incubators were supplemented with either KCl or Pozzolana. Note that the WIPP Surface and the Minus data are the same as shown in Figure 2 above.

The PCA plots from the Study I of the two supplemented sources of radiation compared to the below background minus radiation treatment are shown in **Figure 5**. It is evident that male mosquitos are responding to the two supplemental radiation sources with PC-1 as the most important separator. When documenting the genes that were differentially expressed between the supplemental radiation sources, only the KCl treatment had DEGs associated with any GO terms, with proteolysis genes being the most common genes down-regulated [31]. Interestingly, proteolysis genes in the KCl-supplemented treatment were also downregulated in the underground when compared to the Surface control, indicating that KCl functions as an appropriate radiation source to mimic natural radiation, as we’ve been proposing over the years [10, 11,12].

**Figure 5.**
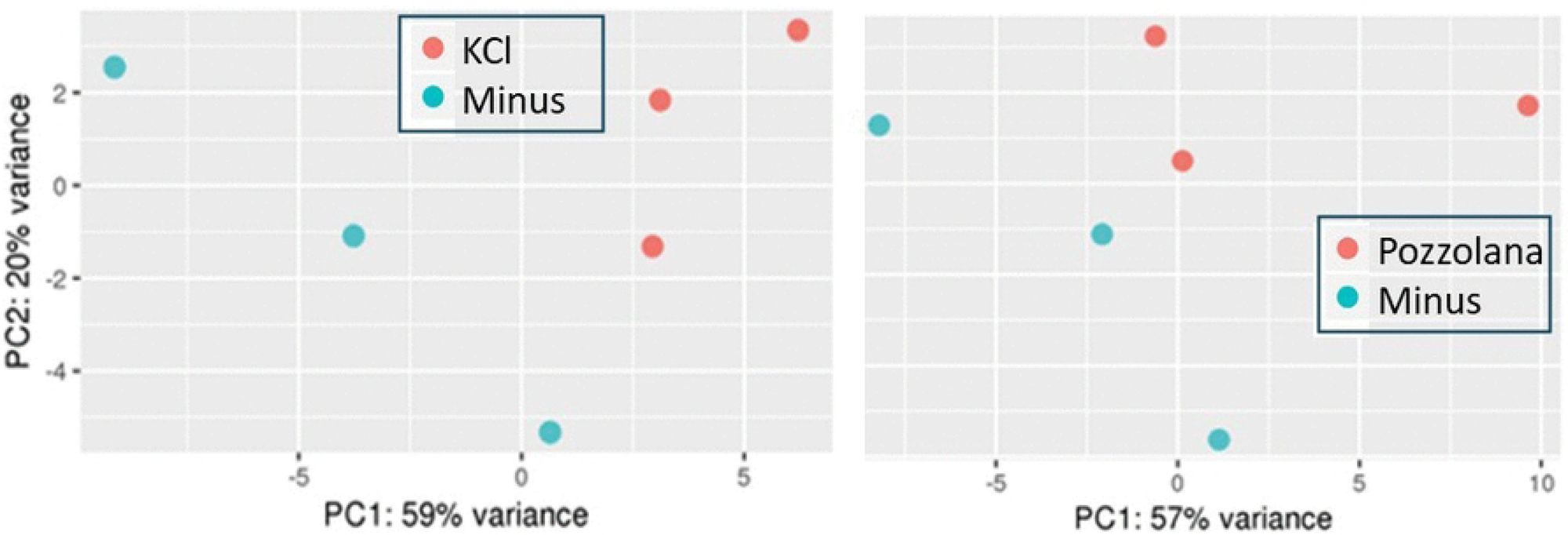
Azenta PCA of the transcriptomes from Study I comparing amendment of two natural sources of radiation compared to the below background radiation-minus treatment.

## Conclusions

Both the studies showed *Ae. aegypti* having a higher mortality rate when male mosquitos were grown in the absence of normal levels of radiation. This trend showing a fitness cost to growth in sub-background levels of radiation is consistent with our previous work with bacteria and mammalian cells [11,12] as well as other research groups studying *Drosophila* [3,4], mammalian cells [5] and yeast [1,7]. However, mating success and concomitant egg laying from females mating with these males were not affected using the methods reported here.

The transcriptome responses to sub-background radiation were different between the two years, with less response documented during the incubation when there were lower levels of mortality. The data suggest that in *Ae. aegypti* a secondary stressor was needed in order to accentuate the gene expression differences due to the stress of reduced radiation.

In spite of very similar levels of radiation, two natural radiation sources (KCl and pozzolana) evoked different gene expression patterns indicating that the source of radiation, that is the quality of the radiation, not just the quantity, is an important factor to consider in radiation studies.

## Acknowledgements

We thank Los Alamos National Laboratory’s Shawn Otto and Jon Davis for their technical and safety support in the Waste Isolation Pilot Plant underground.

